# PENGUINN: Precise Exploration of Nuclear G-quadruplexes Using Interpretable Neural Networks

**DOI:** 10.1101/2020.06.02.129072

**Authors:** Eva Klimentova, Jakub Polacek, Petr Simecek, Panagiotis Alexiou

## Abstract

G-quadruplexes (G4s) are a class of stable structural nucleic acid motifs that are known to play a role in a wide spectrum of genomic functions, such as DNA replication and transcription. The classical understanding of G4 structure points to four variable length guanine strands joined by variable length stretches of other nucleotides. Experiments using G4 immunoprecipitation and sequencing experiments have produced a high number of highly probable G4 forming genomic sequences. The expense and technical difficulty of experimental techniques highlights the need for computational approaches of G4 identification. Here, we present PENGUINN, a machine learning method based on Convolutional Neural Networks, that learns the characteristics of G4 sequences and accurately predicts G4s outperforming the state-of-the-art. We provide both a standalone implementation of the trained model, and a web application that can be used to evaluate sequences for their G4 potential.

## Introduction

G-quadruplexes (G4s) are stable secondary structures of nucleic acids that occur when quartets of guanines are stabilized by a monovalent cation (Gellert, Lipsett, and Davies 1962) and form a characteristic layered structure (Sen and Gilbert 1988) (Figure 1a). G4s are known to play important roles in several biological processes, such as DNA replication, damage response, RNA transcription and processing, transcriptional and translational regulation and others (Spiegel, Adhikari, and Balasubramanian 2020). Owing to their importance as modulators of genomic function, G4s have been studied extensively, and several attempts have been made to model their structure in a predictive manner and several experimental methods for their identification have been developed (Puig Lombardi and Londoño-Vallejo 2020).

**Figure 1:**
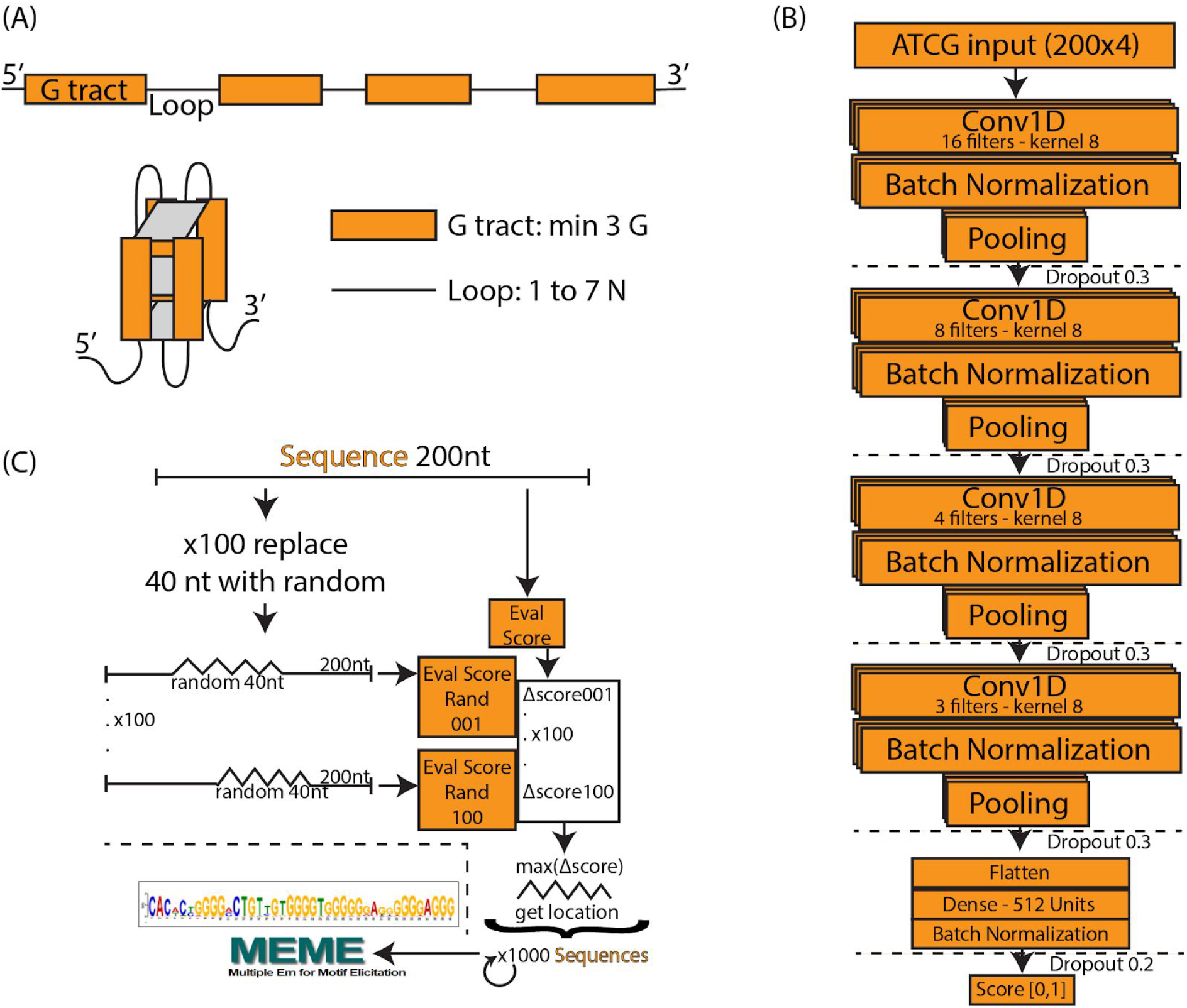
a) Schematic of a typical G-quadruplex structure consisting of four G tracts with a minimum length of 3, connected by non specific loops. b) PENGUINN convolutional neural network model. c) identification of G-quadruplex subsequences via randomized mutation.

Early methods of G4 prediction were focused on the identification of a consensus motif, using a regular expression matching approach, often complemented by involved scoring calculations. An example of such methods is Quadparser (Huppert 2005). It was not until a high-throughput sequencing method for genome wide identification of G4s (G4-Seq) was established (Chambers et al. 2015) that we started understanding how common G4s were, and how hard it is to accurately predict their genomic location. Out of over 700 thousand G4s identified in the human genome by high-throughput sequencing, approximately 450 thousand were not predictable by computational methods at the time. The incentive for the improvement of G4 prediction computational methods and a dataset that would allow us to do so, became evident. A second wave of computational methods attempted to predict G4 locations after the publication of this dataset. Among the most accurate and still functional methods are G4Hunter (Bedrat, Lacroix, and Mergny 2016), which expands the regular expression methods by scoring on G enrichment, Pqsfinder, which focuses on allowing customization for non-canonical G4s and was trained on the G4-Seq dataset (Hon et al. 2017), as well as Quadron (Sahakyan et al. 2017), a Machine Learning method trained on the G4-Seq dataset, utilizing the tree based Gradient Boosting Machine approach.

This second generation of G4 identification methods utilizes Machine Learning to classify sequences based on their G4 forming potential. Generally, Machine Learning (ML) describes the field of computer science that implements mathematical models which enable computers to learn concepts and patterns embedded in data. One of the largest subfields of ML deals with the development of artificial Neural Networks (NNs), which were initially proposed as simplified models of neuronal function (Fitch 1944) and have recently revolutionized the fields of speech recognition and image classification (LeCun, Bengio, and Hinton 2015). The recent breakthrough in the field of NNs involves the utilization of Deep NNs consisting of a large number of neuronal layers. A specific subset of these Deep NNs uses a process known as convolution to learn increasingly complex representations of patterns in raw data. These NNs are called Convolutional Neural Networks (CNNs). An important characteristic of CNNs is their ability to operate on raw data such as images, time-series, DNA/RNA sequences, without the need for complicated feature extraction. The flipside of this ability is their need for large amounts of data. Coupled with the novel availability of high-throughput biological data (Emmert-Streib et al. 2020), Deep NNs are quickly becoming feasible in the field of bioinformatics (Tang et al. 2019). Another important current field of research is the interpretation of Deep NN models, which are often seen as ‘black boxes’ due to their complexity. Convolutional Neural Networks for G4 prediction were implemented in the method G4detector (Barshai and Orenstein 2019).

Here, we present PENGUINN, a CNN based approach for the identification of G4s from raw DNA sequence data, trained on G4-Seq high throughput human data. We establish that PENGUINN outperforms the state-of-the-art in a high background testing set that simulate high genomic variation, and interpret aspects of the learned model, validating its learning against known characteristics of G4 sequences. All data, training scheme, trained models, and functional code can be found at https://github.com/ML-Bioinfo-CEITEC/penguinn. An easy to use Web Application that can run the trained model for user submitted sequences in real time is also made available at https://ml-bioinfo-ceitec.github.io/penguinn/.

## Materials and Methods

### Training and Evaluation Datasets

Our dataset was generated from high-throughput sequencing of DNA G-quadruplexes from the human genome. We used genomic coordinates obtained from a G4-seq experiment (Chambers et al. 2015) (GEO: GSE63874). The coordinates were in three separate sets analogous to three different stabilizers - K+, PDS and K+ together with PDS. Using the original bed files and the hg19 genome annotation we extracted DNA sequences using bedtools (Quinlan and Hall 2010). All sequences which were longer than 200 nt were centred and cut to the length of 200 nt. Shorter sequences were randomly padded from both sides with Ns to become 200 nt long. We referred to this adjusted set as the positive set of the classification problem. For every sequence in our positive set we created a negative sequence of the same length from a random coordinate from hg19 that did not overlap with any of the coordinates from the positive set. Sequences shorter than 200 nt were again randomly padded with Ns. The sequences thus obtained formed a negative set.

We randomly selected 300k sequences (150k positive and 150k negative) from the samples as a training set with pos:neg ratio 1 (the 1:1 dataset). We also randomly selected 300k sequences to create the pos:neg training datasets of 1:9, 1:99 and 1:999, containing 30k, 3k and 300 positives respectively. These datasets correspond to 50%, 10%, 1% and 0.1% positive admixtures respectively.

We selected four sets consisting of 100k sequences each as our final evaluation sets, never seen during any training step. The individual test sets have the same pos:neg ratio as the training sets - 1:1, 1:9, 1:99 and 1:999.

### Training Scheme

We utilized a Convolutional Neural Network consisting of four convolution layers with kernels of size 8 and 16, 8, 4 and 3 filters respectively. The output of each convolutional layer goes through a batch normalization layer, max-pooling layer and dropout layer with the dropout rate 0.3. The output of the last layer is flattened and goes through a densely-connected layer with ReLU activation function. The last layer is formed of a single neuron with a sigmoid activation function, which assigns to each input DNA sequence a probability of having a G4 structure. Our model was implemented in Python using the Keras library with Tensorflow backend. We used Adam optimizer with β_1_ = 0.9 and β_2_ = 0.99, the learning rate was set to 0.001. The loss function was binary crossentropy, metrics accuracy. The model was trained over 15 epochs, the chosen batch size was 32. Figure 1B outlines the architecture of the network.

### Evaluation Scheme

We evaluated our model against five other state of the art methods. First, we tested against the widely used regular expression ‘(G{3,}[ATGCN]{1,7}){3,}G{3,}’, also used by a tool called Quadparser (Huppert 2005). We implemented the regular expression in python, returning a boolean expression dependent on the presence of a match in the presented sequence. The remaining three methods that were developed for the scoring of G4 forming potential are G4Hunter (Bedrat, Lacroix, and Mergny 2016), Quadron (Sahakyan et al. 2017) and Pqsfinder (Hon et al. 2017). We re-implemented G4Hunter in python as the available code did not appear functional (code available at our repository). We used a window of size 25 nucleotides as proposed in the original paper, and a score threshold 0 to see all putative G4s. For every input sequence, the output of our implementation is the highest score of all subsequences. If no G4 has been found, the output score is 0. We ran Quadron with the default parameters. For every input sequence, we considered only scores assigned to the plus strand and we took the maximum of all scored G4s present in the sequence. If no score has been assigned, the output score was zero. For testing Pqsfinder was used following command: ‘pqsfinder(sequence, strand=‘+’, overlapping=TRUE, verbose=FALSE)’, as an output we took the highest scoring G4, if none has been found, the score was set to 0. Lastly, we compared our model to another machine learning model G4detector (Barshai and Orenstein 2019). We ran it in the testing mode using three available models trained on random negatives and positives with K^+^, PDS and K^+^ + PDS stabilizers.

### Interpretation of Model

In order to interpret the model, we have attempted to isolate the particular features detected by the model which weigh the most for its decision making. For each positive testing sample we have produced 100 sequences of the same length containing a random stretch of 40 nucleotides for every possible position. For each such sequence we re-evaluate and calculate the average degree of score change for the subsequence. The subsequence that produces the largest drop is marked as the ‘most important’ and extracted (Fig. 1c).

### Code availability and web application

PENGUINN was developed in Python. All code accompanied by the trained models, all training data and the installation instructions can be found at https://github.com/ML-Bioinfo-CEITEC/penguinn. Moreover, we have converted the trained PENGUINN Keras model into TensorflowJS and developed a simple web application, available at https://ml-bioinfo-ceitec.github.io/penguinn/. The web application code can be found in the gh-pages branch of the PENGUINN GitHub repository.

## Results

Scanning across genomic regions for a specific and relatively rare structural element is a task that involves a heavy class imbalance, since the background sequence will heavily outnumber the target element by orders of magnitude. It is hard to know the exact prevalence of G4s in the genome or at least the areas of the genome one would scan, but the more imbalance datasets should approximate a realistic ratio more closely. A rough estimate could come from equally dividing the approximately 700 thousand known G4s over the 6 billion bases of the human genome, giving us an approximate ratio of a G4 located every 8 thousand base pairs. For this reason, we have produced four datasets with increasing positive to negative ratio (pos:neg) by one order of magnitude each time. Starting from the highly unrealistic 1:1 dataset with equally balanced classes (50%) and then going up to 1:9 (10%), 1:99 (1%), and 1:999 (0.1%) ratio datasets. We have also acquired a dataset (Puig Lombardi and Londoño-Vallejo 2020) with high class imbalance in the opposite direction consisting of 298 positives and 94 negatives (3:1 dataset).

Initially, we explored the possibility of training models in equally imbalanced datasets and then using them to improve prediction accuracy. However, we could not see any measurable improvement for training with a matching pos:neg mixture when considering the area under the precision sensitivity curve for our models, or when using an iterative negative selection technique that previously showed improvement in a different genomic classification task (Georgakilas et al., n.d.)(Supp. Fig. S1). Since there does not appear to exist a major difference in performance between these models, we have elected to use the model trained on 1:1 as our main trained model. Plotting the F1 score against the prediction score of our method for each testing dataset (Supp. Fig S2) we have identified two score values that we proposed as score thresholds for our method (precise: 0.85, sensitive: 0.5). Users are allowed to set their own cut-off threshold for their results depending on their needs, but having proposed score thresholds helps new users guide their decisions to more meaningful thresholds. For clarity of presentation, on all evaluations against state of the art we will designate these thresholds as PENGUINN(s) for the sensitive, and PENGUINN(p) for the precise threshold.

A commonly used method for G4 identification is the use of a sequence pattern, also called a regular expression, consisting of up to four stretches of Gs with a minimum length of three, spaced by random nucleotide sequences with a maximum length of seven (for exact expression see Materials and Methods). This method was first proposed over 15 years ago (Huppert 2005) and has been commonly used since then. We have directly compared PENGUINN to this regular expression in all our testing datasets. Since the regular expression cannot return a score and will only produce a binary result, it is not possible to produce a ROC curve or similar metric across scores. Our models outperformed the G4 regular expression in all datasets with increasing difference as datasets became more negative heavy (Table 1, Figure 2). Despite being a widely used way of identification for G4s, the regular expression lacks both precision and sensitivity compared to more elaborate methods such as PENGUINN.

**Table 1:**
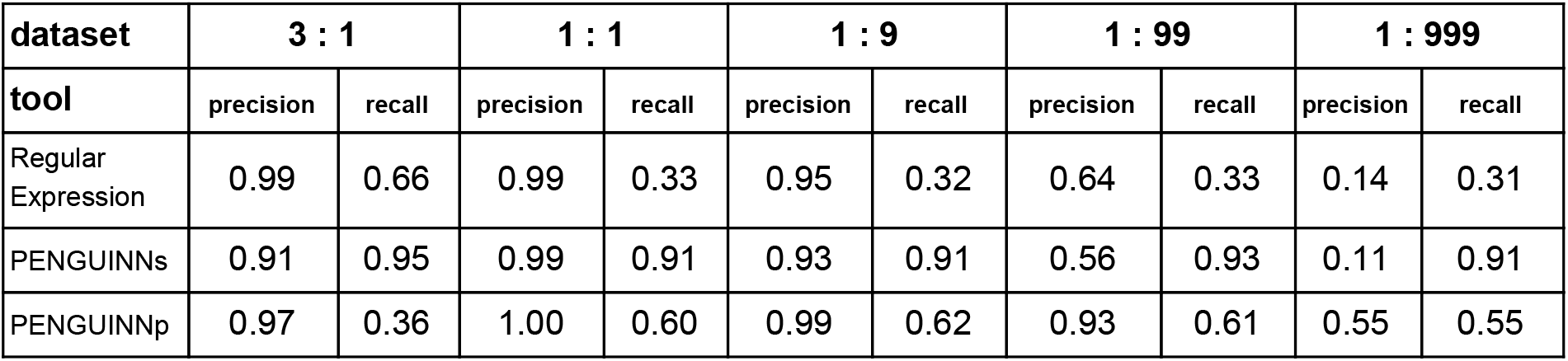
Precision and Recall values for static score prediction of Regular Expression, PENGUINNs (sensitive) and PENGUINNp (precise) on a scale of imbalanced datasets.

**Figure 2:**
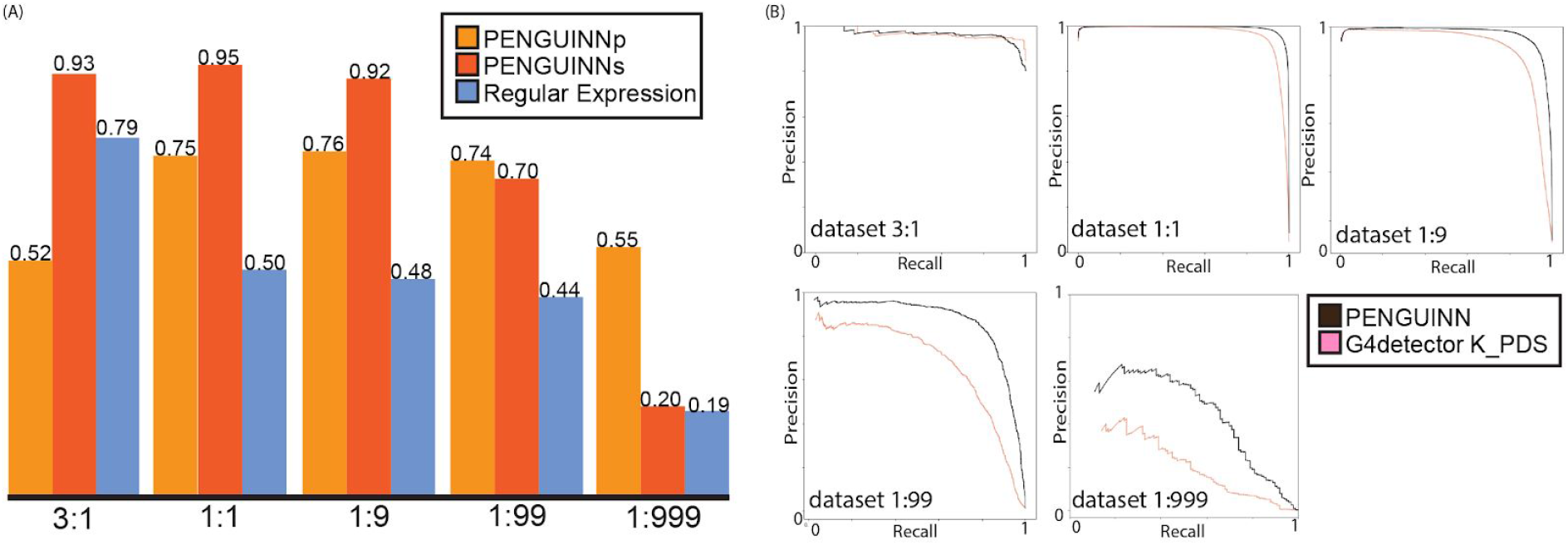
(A) F1 score for PENGUINNp (precise), PENGUINNs (sensitive) and Regular Expression with datasets of different pos:neg ratio. (B) Precision-Recall curve comparison of PENGUINN and best performing state of the art method G4detector K_PDS in datasets of different pos:neg ratio.

We proceeded to evaluate our method against 4 other state-of-the-art methods on the same benchmark datasets. We compared their performance across the whole range of prediction scores using the precision-recall area under curve for each evaluation dataset. There is an evident trend of quickly diminishing performance as datasets become more realistic in ratios with more negatives. Our method also loses performance under these circumstances, but at a much slower rate, pointing at a comparative improvement when used for realistic highly imbalanced datasets (Table 2). We have selected the best performing state of the art method for direct comparison using detailed precision-recall curves for each dataset (Figure 2B). It becomes evident that as the class imbalance increases, both methods lose performance, but PENGUINN manages to retain a higher level of precision/sensitivity even at highly imbalanced datasets. Comparison with all other state of the art programs shows similar patterns as ratios become more realistic. (Supplementary Figures 3-6)

**Table 2:** Area under the Precision-Recall curve for PENGUINN and 4 state-of-the-art programs. Underlined are the best performances for each evaluation dataset.

| Dataset | 3 : 1 | 1 : 1 | 1 : 9 | 1 : 99 | 1 : 999 |
| --- | --- | --- | --- | --- | --- |
| PENGUINN | 0.96 | <u>1.00</u> | <u>0.97</u> | <u>0.87</u> | <u>0.43</u> |
| G4detector K PDS | 0.97 | 0.98 | 0.91 | 0.67 | 0.21 |
| G4detector PDS | 0.94 | 0.98 | 0.91 | 0.63 | 0.21 |
| G4detector K | 0.94 | 0.98 | 0.89 | 0.58 | 0.16 |
| G4Hunter | 0.97 | 0.97 | 0.86 | 0.54 | 0.13 |
| Quadron | 0.97 | 0.83 | 0.67 | 0.50 | 0.25 |
| PQSfinder | <u>0.98</u> | 0.95 | 0.86 | 0.58 | 0.15 |

Deep Learning models consist of complex networks of neurons that abstract information in hard-to-interpret ways. Often these models are considered uninterpretable black boxes. However, this can lead to pitfalls such as learning data artifacts instead of real signal. To control against such pitfalls, we identified what our model considered the most important 40 nucleotide subsequence for 1000 samples, using a randomized permutation approach. We sorted by the degree of change when randomized and used the top 1000 of these sequences to identify enriched motifs using Multiple Em for Motif Elicitation (MEME) (Bailey et al. 2006). The top motif extracted (Fig 1c) is indeed a motif containing several G-tracts which confirms the known theory of G4 formation and demonstrates that our model did not primarily learn some artifact such as padding length. The motif produced by MEME is an aggregate of several similar motifs found in these sequences, and as such is not expected to appear as an exact ‘consensus’ G4.

Paradoxically, although elaborate ML models, such as PENGUINN, vastly outperform the simple regular expression search for G4s, its use persists to date. Beyond the familiarity of the method, we believe that any technical obstacle, however trivial, will deter non-technical users from using other methods. As such, we decided to create a straightforward web application that uses our best trained model to evaluate user submitted sequences in real time. The web application can be found here: https://ml-bioinfo-ceitec.github.io/penguinn/. The user can input a single sequence, a fasta formatted input, or several sequences in multiple lines. The sequences will be evaluated, and a score along with threshold evaluation returned.

## Discussion

In this study we present PENGUINN, a convolutional neural network based method that outperforms state of the art methods in the identification of nuclear G4s in highly imbalanced datasets. PENGUINN is more robust than other methods when the pos:neg ratio increases by several orders of magnitude. However, there is still space for improvement in the prediction. We believe that a more elaborate modelling of the real variation of the background genome could benefit predictive methods of this type. Such undertaking is beyond the scope of this study.

Beyond the development of a highly effective predictive model, we have explored the interpretation of what the model has learned. As expected, the model identified regions of high G content as better potential targets, and has scored very highly regions showing periodic G stretches, a structural feature known to define G4s. Convolutional Neural Networks are notorious for being hard to interpret, as deeper network layers further abstract information from the first layers. We believe that interpreting the network to the extent that we can conceptualize the type of sequences it has learned to identify is an important step for genomic sequence deep learning studies.

To allow for easier adoption of our method, we have developed both a standalone version and a web application that can be used without any knowledge of programming. The repository https://gitlab.com/RBP_Bioinformatics/penguinn contains all models, data, and thorough installation and usage tutorials. The web application can accept sequences ranging from 40nt up to hundreds of nts. For sequences smaller than 200nt, our method will pad the sequence with Ns randomly on each side. This may create a variation in scores for really short input sequences. For sequences larger than 200nt, our method will extract 200nt around the midpoint of the sequence. This means that the whole sequence is not evaluated but just the middle 200nt. Users should attempt to preprocess their data as much as possible, centering their potential G4 sequence.

In conclusion, PENGUINN is a powerful method, based on cutting edge Deep Learning architecture, that increasingly outperforms the state of the art in classifying G4s in more realistic highly imbalanced datasets. Despite the sophistication of the method, we have developed a simple web application to assist users coming from non-bioinformatic backgrounds to use the method. We also provide all training and testing datasets in an effort to empower researchers to produce better, more accurate methods for realistic highly imbalanced datasets.

