## Supplementary figures and images for "PENGUINN: Precise Exploration of Nuclear G-quadruplexes Using Interpretable Neural Networks"

### Supplemental Figure S1

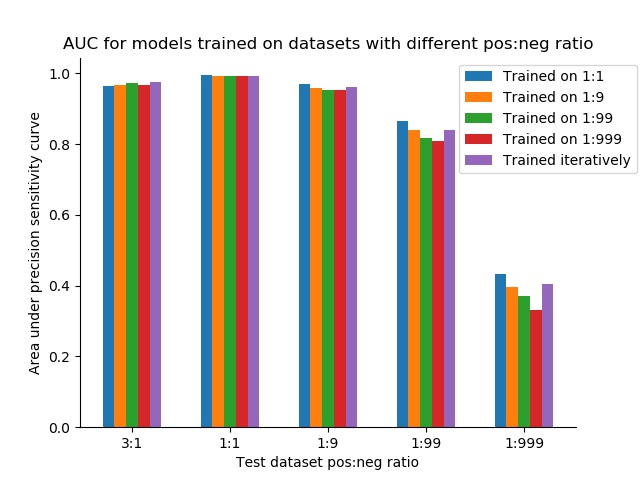

### Supplemental Figure S2

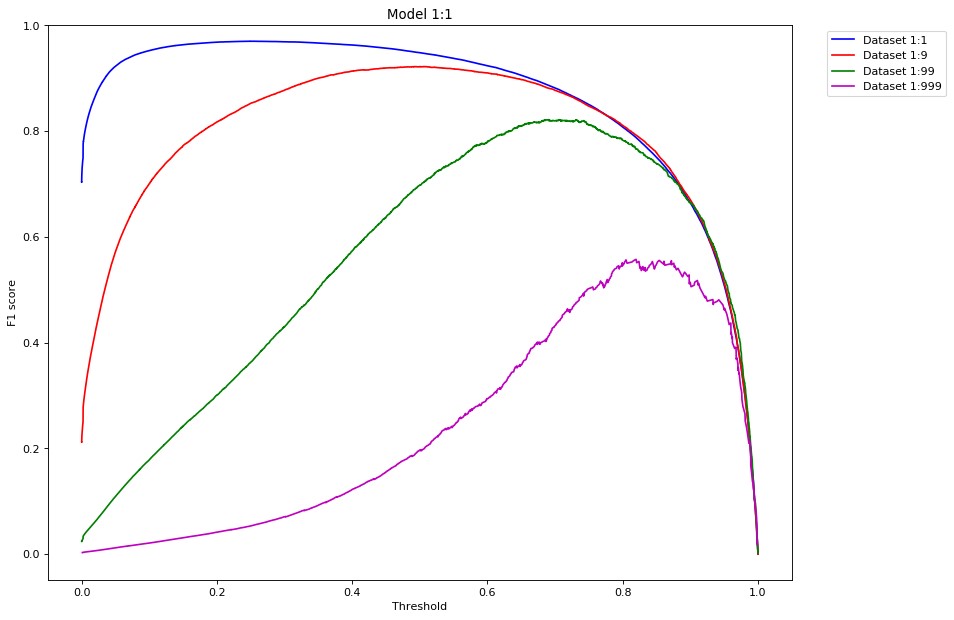

### Supplemental Figure S3

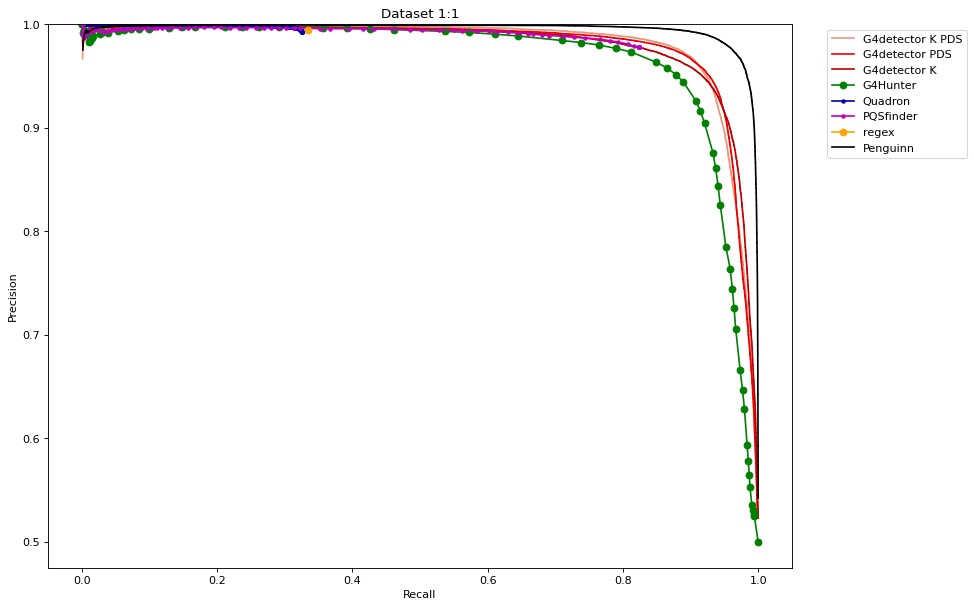

### Supplemental Figure S4

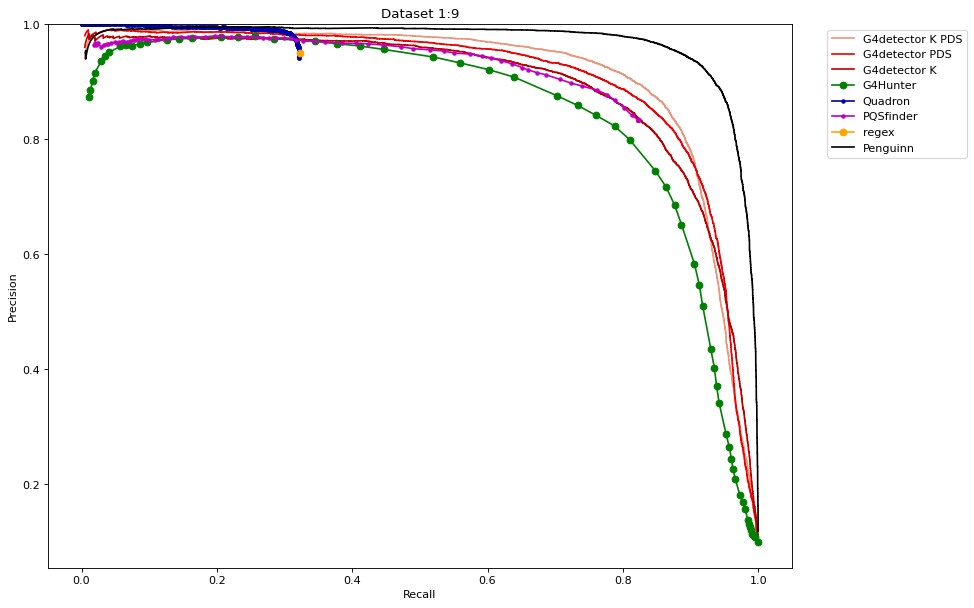

### Supplemental Figure S5

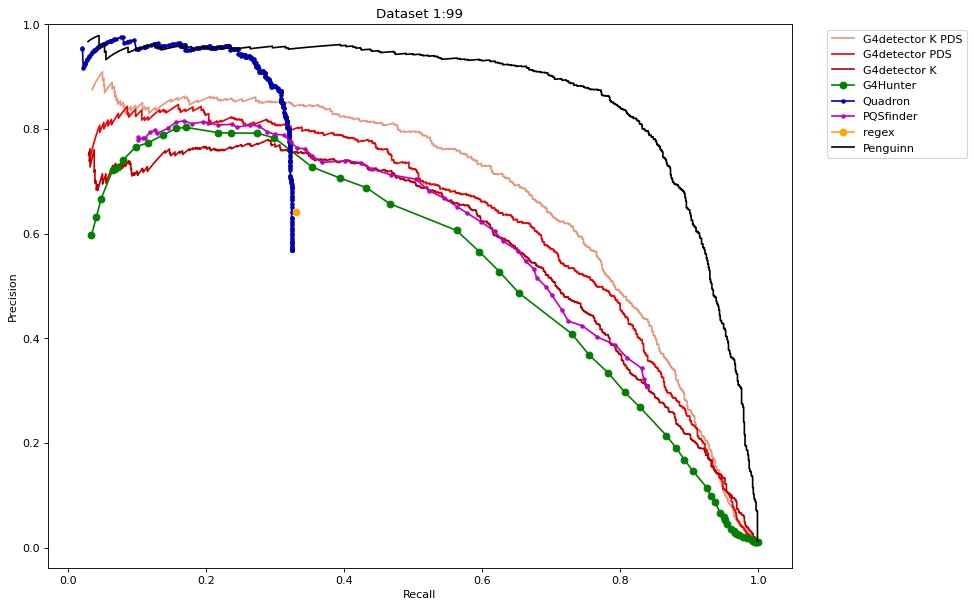

### Supplemental Figure S6

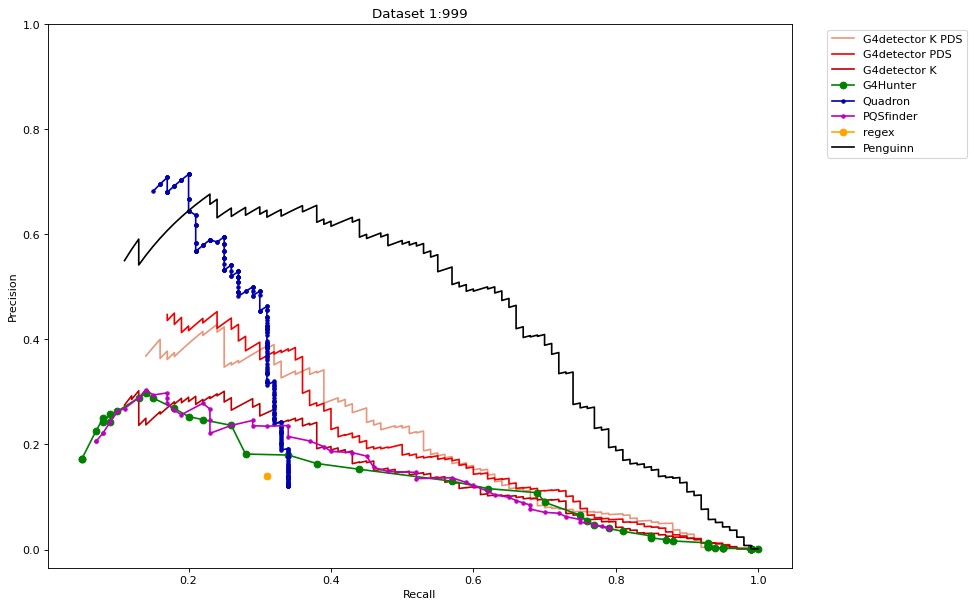
